# Species delimitation in the presence of strong incomplete lineage sorting and hybridization: lessons from *Ophioderma* (Ophiuroidea: Echinodermata)

**DOI:** 10.1101/240218

**Authors:** Alexandra Anh-Thu Weber, Sabine Stöhr, Anne Chenuil

**Author notes:** Corresponding author; ORCID: 0000-0002-7980-388X.

## Abstract

Accurate species delimitation is essential to properly assess biodiversity, but also for management and conservation purposes. Yet, it is not always trivial to accurately define species boundaries in closely related species due to incomplete lineage sorting. Additional difficulties may be caused by hybridization, now evidenced as a frequent phenomenon. The brittle star cryptic species complex *Ophioderma longicauda* encompasses six mitochondrial lineages, including broadcast spawners and internal brooders, yet the actual species boundaries are unknown. Here, we combined three methods to delimit species in the *Ophioderma longicauda* complex and to infer its divergence history: i) unsupervised species discovery based on multilocus genotypes; ii) divergence time estimation using the multi-species coalescent; iii) divergence scenario testing (including gene flow) using Approximate Bayesian Computation (ABC) methods. 30 sequence markers (transcriptome-based, mitochondrial or non-coding) for 89 *O. longicauda* and outgroup individuals were used. First, multivariate analyses revealed six genetic clusters, which globally corresponded to the mitochondrial lineages, yet with many exceptions, suggesting ancient hybridization events and challenging traditional mitochondrial barcoding approaches. Second, multi-species coalescent-based analyses confirmed the occurrence of six species and provided divergence time estimates, but the sole use of this method failed to accurately delimit species, highlighting the power of multilocus genotype clustering to delimit recently diverged species. Finally, Approximate Bayesian Computation showed that the most likely scenario involves hybridization between brooders and broadcasters. Our study shows that despite strong incomplete lineage sorting and past hybridization, accurate species delimitation in *Ophioderma* was possible using a combination of complementary methods. We propose that these methods, especially multilocus genotype clustering, may be useful to resolve other complex speciation histories.

**Highlights:**

- Multivariate analysis was used for species delimitation
- Six *Ophioderma* species were delimited using nuclear and mitochondrial data
- *Ophioderma* speciation history is complex and included hybridization
- Mitochondrial and nuclear histories differed, challenging barcoding approaches
- We propose that using multilocus genotypes can resolve complex speciation histories

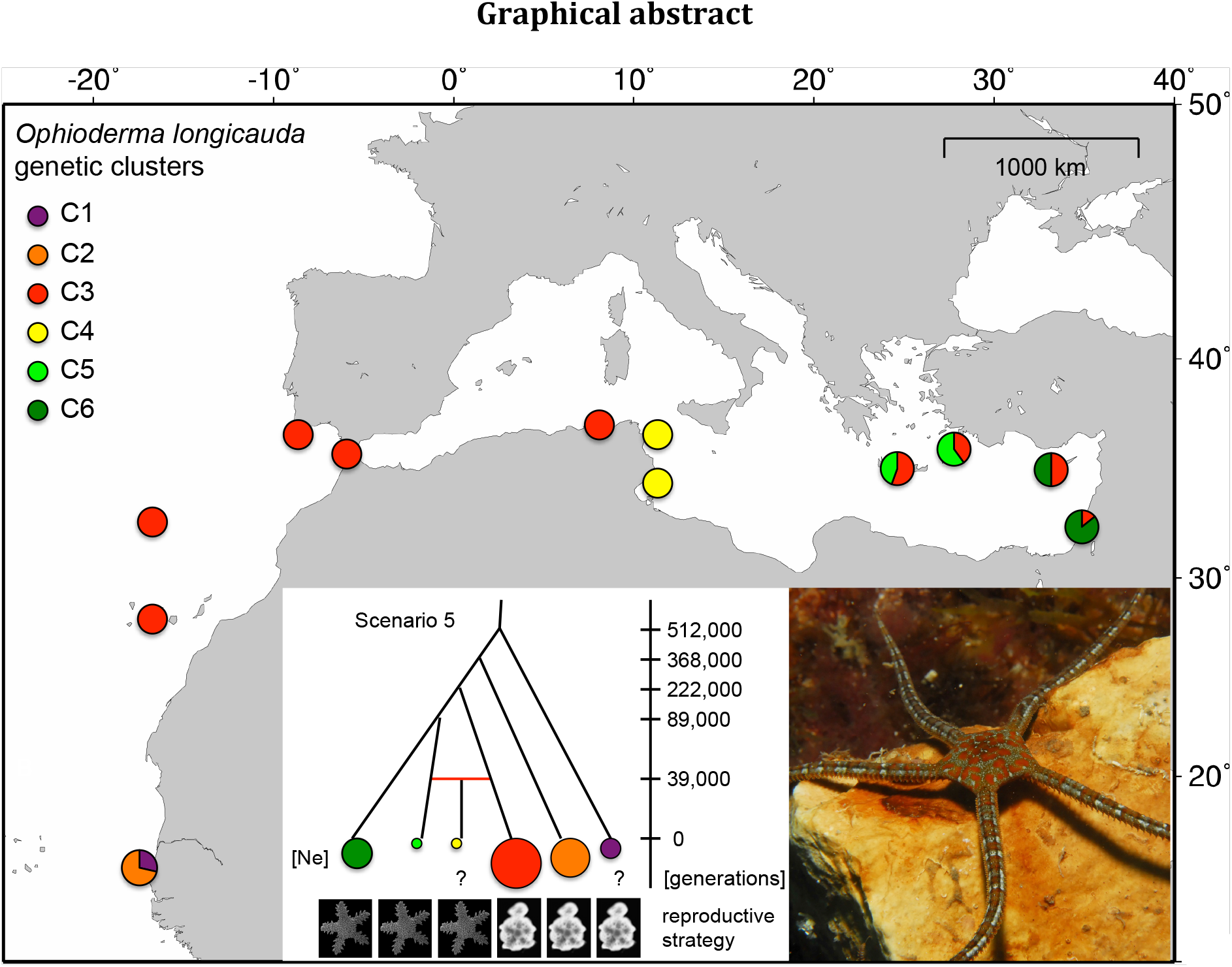

## 1. Introduction

Accurate species delimitation and description is essential to properly assess biodiversity, but also for management and conservation purposes (Agapow et al., 2004; De Queiroz, 2007). Historically and still nowadays, the vast majority of species are delimited using morphological characters, based on descriptions of type specimens for each nominal species, uniquely identified by Latinized names in the binominal nomenclature codified first by Linnaeus in the 18th century (for zoology (Linnaeus, 1758)). However, genetically isolated groups of morphologically indistinguishable individuals were detected in many nominal species during the last decades, owing to the use of genetic markers in population genetic studies (Bickford et al., 2007; Knowlton, 1993; Pfenninger and Schwenk, 2007). Such groups are often called cryptic species (Struck et al., 2018), which are widely spread and homogeneously distributed across the metazoan biodiversity (Pfenninger and Schwenk, 2007). The occurrence of cryptic species can be explained with several main factors: i) recent species divergence (i.e. morphological differences may not have evolved yet); ii) parallelism or convergence (i.e. cryptic species are not closely related but morphologically similar due to selection pressures); iii) morphological stasis (i.e. decreased morphological disparity compared to genetic divergence) (Struck et al., 2018)). Cryptic species were first identified by diagnostic codominant markers such as allozymes (e.g. Knowlton, 1993) or by single mitochondrial markers. Even presently, they are rarely identified using a combination of several molecular markers and phenotypic data (Struck et al., 2018).

Due to their lower effective size, genetic markers from haploid genomes are more affected by genetic drift and thus reach reciprocal monophyly (i.e. alleles of distinct species form separate monophyletic groups) more rapidly than markers from nuclear genomes after species divergence. This explains their power to detect recently diverged cryptic species and their wide use for biodiversity barcoding. However, absence of gene flow among groups of individuals cannot be established on the basis of markers from single haploid genomes since past bottlenecks or selective sweeps may generate patterns of divergent groups of closely related haplotypes within a panmictic entity (i.e. a group of randomly mating individuals). In addition, mitochondrial markers only reflect the history of maternal lineages, which can be significantly different from the species history if males and females display different dispersal behaviors. Finally, past hybridization events can be misleading for species identification based on mitochondrial barcodes, as mitochondrial lineages can be incongruent with the species history (Currat et al., 2008; Melo-Ferreira et al., 2014).

Finding independent markers confirming mitochondrial divergence may be difficult for recently diverged species, especially in non-model organisms with scarce to non-existing available genomic data. The most intuitive and popular approach for cryptic species delimitation consists of finding independent markers displaying reciprocal monophyly that are congruently associated within individuals (in linkage disequilibrium) (De Queiroz, 2007; Mkare et al., 2017). However, confirming absence of gene flow should not rely on finding reciprocal monophyly in independent markers (Zhang et al., 2011) for the following reasons: i) Recently diverged species, especially when their reproductive isolation is conferred by a single locus or few loci, rarely display reciprocally monophyletic markers, as most loci are either monomorphic or display shared polymorphism among species (i.e. incomplete lineage sorting). ii) Recently diverged species may display diagnostic markers but not reciprocal monophyly when diagnosticity resulted from genetic drift (allele loss) but mutation events were not sufficient to create a pattern of reciprocal monophyly. iii) There are other means to establish absence of gene flow using a set of independent genetic markers (e.g. the diagnosticity of a single Mendelian marker is sufficient, and see approaches based on multilocus genotype developed below).

Various methods (reviewed in (Carstens et al., 2013) propose to delimit species using nucleotide sequences from several markers. Some of these methods are based on divergence levels or consider ratio of within-group and between-group diversity to delimit species (e.g. the automatic barcoding gap discovery ABGD, (Puillandre et al., 2012). Although they provide powerful tools to propose primary species hypotheses for large datasets (Ratnasingham and Hebert, 2007), such approaches cannot prove the absence of gene flow and often depend on arbitrary thresholds or assumptions such as constant effective sizes among ancestor and daughter species. The multispecies coalescent theory (Fujita et al., 2012; Yang and Rannala, 2010) provides a statistical framework to model the coalescence of multiple markers during genetic isolation of groups of individuals. Bayesian methods applied to the multispecies coalescent allow establishing the probability of species partitions and phylogenies for a sample of allele sequences from various individuals (Rannala, 2015; Yang and Rannala, 2010) and the most recent development of this method, implemented in the software BPP (Yang, 2015), allows the joint inference of species (or lineage) delimitation and species phylogeny (Rannala and Yang, 2017; Yang and Rannala, 2014). However, despite these qualities and being able to handle some degree of Incomplete Lineage Sorting (ILS) (Carstens et al., 2007), the efficiency of these methods depend for a large part on the accumulation of mutations since species separation, and thus can only delimit entities isolated for long enough (Rannala, 2015). By contrast, methods based on multilocus genotypes (e.g. Structure (Falush et al., 2003), Structurama (Huelsenbeck et al., 2011), DAPC (Jombart et al., 2010)), although they are rarely used for species delimitation, can potentially reveal absence of gene flow after a single generation as they do not rely on information on allele relationships, like sequences, but use multilocus genotypes for each individual. In addition, they provide a fast and unbiased way of finding genetically separated entities, as they do not rely on *a priori* knowledge on individual grouping.

Even though much progress has been made in species tree estimation methods, the use of species trees implies that speciation is represented as a dichotomic process. Yet, there is increasing evidence for the role of hybridization and reticulate evolution in or after speciation (Abbott et al., 2013, 2016; Lamichhaney et al., 2017; Meier et al., 2017). Current species discovery and delimitation methods do not allow for testing such cases, but Approximate Bayesian Computation (ABC) provides powerful methods allowing to do so (Csilléry et al., 2010; Lopes and Beaumont, 2010), with the simultaneous testing of several divergence scenarios (with or without hybridization) and estimation of demographic parameters (e.g. divergence times, effective population sizes, migration rates). These methods are computationally efficient, as they use simulated datasets for which several summary statistics are compared to the original dataset (instead of likelihood computations). Thus, information rich datasets such as sequence genotypes at tens of loci in a hundred individuals can be exploited using ABC.

Brittle stars (Ophiuroidea) encompass a large number of cryptic species (e.g. (Barboza et al., 2015; Baric and Sturmbauer, 1999; Boissin et al., 2017, 2008; Heimeier et al., 2010; Hoareau et al., 2013; Hunter and Halanych, 2008; Muths et al., 2009, 2006; Naughton et al., 2014; Pérez-Portela et al., 2013; Sponer and Roy, 2002; Stöhr and Muths, 2010; Taboada and Pérez-Portela, 2016). The *Ophioderma longicauda* (Bruzelius, 1805) species complex encompasses six mitochondrial lineages and at least two sympatric biological species with contrasting reproductive strategies, namely the broadcast spawners C3 and the internal brooders C5, differing in reproductive timing, morphology, ecology and genetics (Boissin et al., 2011; Stöhr et al., 2009; Weber et al., 2015, 2014, 2013). Yet, the species relationships among all mitochondrial lineages across the whole *O. longicauda* distribution remain unclear.

We used 30 sequence markers to delimit species and infer divergence history of this complex of cryptic species using a combination of three methods: i) unsupervised species discovery using multilocus genotypes; ii) confirmation of lineage genetic isolation and divergence time estimates using the multi-species coalescent; iii) divergence scenario testing using ABC. We found that combining all three methods that use different properties of the data provides complementary information such as number of species, among species relationships, divergence time, effective population size and gene flow estimates to best represent complex speciation history. Furthermore, the use of multilocus genotypes performed better than the multispecies coalescent to delimit recently genetically isolated clusters.

## 2. Material and Methods

### 2.1. Sampling, DNA extraction and marker development

89 individuals including the six *O. longicauda* mitochondrial lineages (L1-L6) sampled across the whole *O. longicauda* distribution and three outgroup species (*Ophioderma teres* (Lyman, 1860), *Ophioderma cinerea* (Müller & Troschel, 1842) and *Ophioderma phoenia* (H.L. Clark, 1918)) were used in this study (Table S1). *Ophioderma* outgroups were used to estimate divergence times of the *O. longicauda* species complex, the latter occurring in the North-East Atlantic Ocean and in the Mediterranean Sea (Fig. 1). Indeed, the species *O. teres* (Eastern Pacific Ocean), *O. phoenia* and *O.cinerea* (West Atlantic Ocean) are geminate pairs that speciated after the closing of the Isthmus of Panama, so the divergence between these species pairs (*O. teres - O. phoenia* or *O. teres - O.cinerea*) is at least 2.8 Mya (Lessios, 2008). DNA was extracted with MN-Tissue Kit (Macherey-Nagel) using an epimotion robot (Eppendorf) following the protocol of Ribout and Carpentieri (2013), and eluted in 200μl of sterile water. Extracted DNA was diluted 10x in sterile water prior to PCR. We used orthologous genes from transcriptomes of *O. longicauda* C3 and C5, corresponding to mitochondrial lineages L1 and L3, respectively (Weber et al., 2017, 2015) to develop 55 primer pairs to test. The criteria for marker development were: (i) the marker should be polymorphic; (ii) in half of the markers, at least one diagnostic SNP between C3 and C5 should be present; (iii) the length of the PCR product should be 300-400 bp (due to sequencing technology limitations). In addition, seven already existing markers were used, namely a mitochondrial marker (COI), ribosomal markers (ITS1, ITS2) and four EPIC markers, introns i21, i36, i50 and i54b (Chenuil et al., 2010; Gérard et al., 2013; Penant et al., 2013). Of the 55 exon-based markers tested, 16 amplified correctly in each lineage of *O. longicauda*. Furthermore, six out of seven existing markers amplified correctly. Finally, 22 markers were PCR amplified in the 89 specimens.

**Figure 1:**
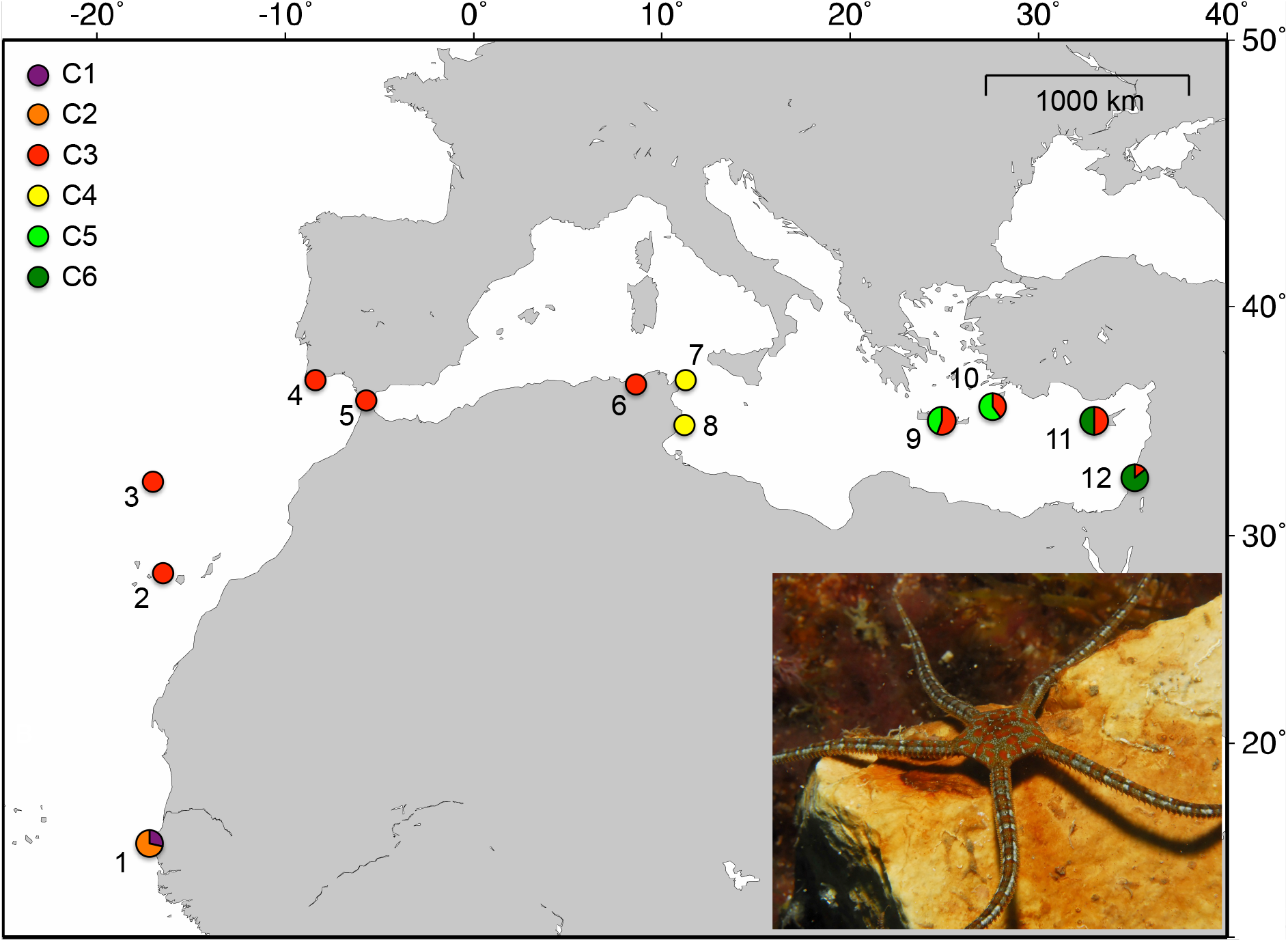
Map of sampling localities with colors corresponding to genetic clusters found with DAPC. 1: Dakar, Senegal. 2: Teneriffe, Canary Islands. 3: Madeira, Portugal. 4: Algarve, Portugal. 5: Ceuta, Spain. 6: Tabarka, Tunisia. 7: Kelibia, Tunisia. 8: Monastir, Tunisia. 9: Agios Pavlos, Crete. 10: Symi island, Greece. 11: Baths of Aphrodite, Cyprus. 12: Ramkine, Beirut and Raoucheh, Lebanon. A photograph of *O. longicauda* C3 from Marseilles, France, is displayed for illustration. Photo credit: Frédéric Zuberer.

### 2.2. Amplicon sequencing and dataset processing

PCR products of different genes belonging to the same individual were pooled and 96 Illumina libraries were constructed. Paired-end (2x 250 pb) sequencing was performed on a MiSeq Sequencing System (Illumina) by the genomic platform Genotoul (www.genotoul.fr). About thirty millions raw reads were obtained after sequencing. Reads were cleaned, assembled and demultiplexed using the program MOTHUR v 1.31.2 (Schloss et al., 2009). On average, between 1000 and 10,000 sequences were obtained per marker and per individual. Then, identical sequences were clustered and the number of reads per sequence and per individual was counted for each marker.

As the number of reads differed greatly between markers (less than 100 reads to more than 1000 reads), applying a fixed threshold to keep final sequences was not possible. In addition, five markers displayed paralogous genes (e.g. more than two sequences with high and similar number of reads displayed). For this reason, selecting the sequence displaying the highest number of reads could lead to incorrectly selecting and clustering paralogous genes. Therefore, for each of the 22 markers, the number of reads obtained for 5-10 individuals was manually checked to determine a threshold to apply to each individual per marker. One (for homozygous individuals) or two sequences (for heterozygous individuals) were kept per individual and per marker when paralogous genes were unambiguously absent, and up to ten sequences per individual and per marker were kept for genes displaying paralogs. Of the 22 markers, three could not be used due to a too low number of reads obtained after sequence cleaning. Finally, a total of 18 genes were obtained, and since five of them displayed paralogs, 30 markers were available for further analyses (Table S2).

### 2.3. Haplotype networks and mitochondrial distances

For each marker, haplotype networks were generated using the median-joining algorithm of Network, version 4.6.1.1 (Bandelt et al., 1999). Kimura 2-parameter (K2P) pairwise distances (Kimura, 1980) among mitochondrial lineages (or among species when considering outgroups) were calculated using MEGA v7 (Kumar et al., 2016). The within-group K2P distances were calculated in the same way.

### 2.4. Species discovery: Principal Component Analysis (PCA) and Discriminant Analysis of Principal Components (DAPC)

In order to determine the number of existing genetic groups without prior knowledge (i.e. mitochondrial lineage or geographic origin), we performed a Discriminant Analysis of Principal Components (DAPC) using the R software package *adegenet* 1.4-1 (Jombart et al., 2010). The DAPC is a clustering method that maximizes the between-group variance while minimizing the within-groups variance. This analysis uses genotypic information for each individual and each locus. Therefore, we converted our sequence data in genotype data using PGDspider v.2.1 (Lischer and Excoffier, 2012). Then, the minimum value of the Bayesian Information Criterion (BIC) was used to determine the optimal number of genetic clusters k, implemented in *adegenet*. After that, DAPC was performed to define the clusters and visualize their relationships. It also provides membership probabilities, i.e. the probabilities for each individual to belong to a particular cluster. Analyses were performed including and excluding the mitochondrial marker COI to infer whether it significantly influenced the genetic clustering. Pairwise F_ST_ were calculated for the six genetic clusters found with the DAPC analysis (see Results) using Genetix (Belkhir et al., 2004). We then performed a PCA on the multilocus diploid genotypes to explore the genetic relationships among individuals without constraint and without a priori knowledge on population membership. We visualized (using colors) the genetic proximity among individuals from the distinct clusters previously identified by the DAPC, but the PCA does not use this information and does not attempt to delimit divergent groups. For this reason, it can suggest incomplete separation or hybridizations between the clusters of individuals visualized by distinct colors.

### 2.5. Lineage confirmation and divergence time estimates: the multi-species coalescent

Since genetic clusters obtained after the first approach from multilocus genotypes appeared as separate genetic entities (see Results) we considered that their relationships could be described by tree-like topologies, possibly assuming gene flow events between some clusters. Based on the genetic clusters found with DAPC, we calculated the average distance between clusters using between-group K2P distances (based on the 30 sequence markers) implemented in MEGA 6.0.5. Then, we reconstructed a phylogenetic tree based on the K2P distances between clusters, using the distance-based Neighbor-Joining method, to define a starting tree for multi-species coalescent based analyses (Fig. 2, scenario 1). Joint Bayesian species delimitation and species tree estimation was conducted using the program BPP v3.3 (analysis A11; (Rannala and Yang, 2017; Yang, 2015)). The method uses the multispecies coalescent model to compare different models of species (or lineage) delimitation (Rannala 2015; Sukumaran & Knowles 2017) and species (or lineage) phylogeny in a Bayesian framework, accounting for incomplete lineage sorting due to ancestral polymorphism and gene tree-species tree conflicts (Rannala and Yang, 2013; Yang and Rannala, 2014, 2010). The population size parameters (θs) are assigned the gamma prior G(2, 1000), with mean 2/2000 = 0.001. The divergence time at the root of the species tree (τ0) is assigned the gamma prior G(2, 100), while the other divergence time parameters are assigned the Dirichlet prior (Yang & Rannala, 2010: equation 2). After 100,000 burnin iterations, 500,000 MCMC samples were recorded with a sample frequency of 100. Each analysis was run three times to confirm consistency between runs. Analyses were run using the full dataset and species were defined using the clusters found with DAPC.

**Figure 2:**
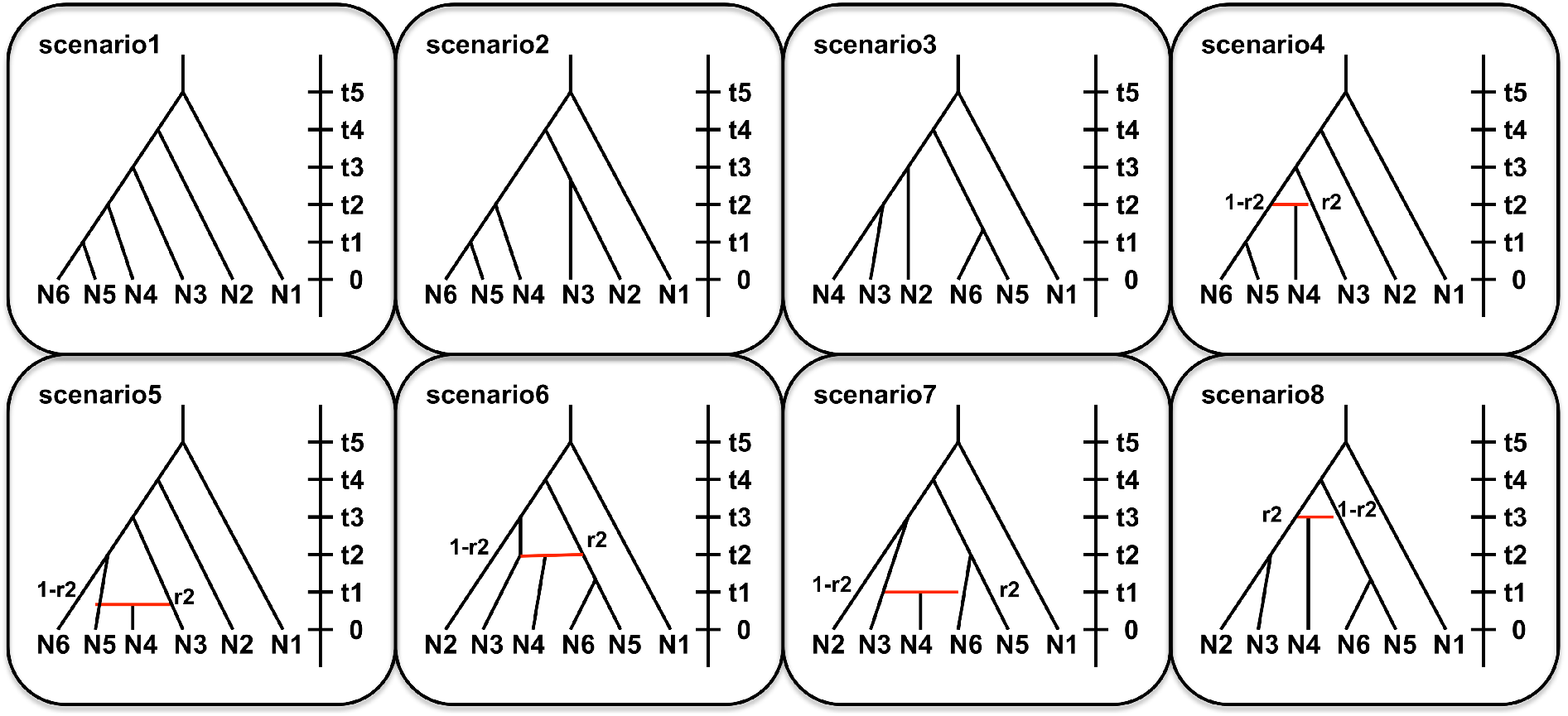
Eight divergence scenarios tested to infer evolutionary history of *O. longicauda* species complex divergence.

As it was recently suggested that ‘species discovery’ methods may eventually not be necessary due to improving of algorithms and computational power (Rannala, 2015), we tested the accuracy of BPP alone to discover and delimit species, using a subset of our dataset for computational purposes. We used all C3 (9 individuals) & C5 (11 individuals) specimens from Greece, known to represent two sympatric biological species (brooding and broadcast spawning individuals; (Weber et al., 2015, 2014)). A DAPC was first performed on this sub-dataset. Then, each individual was set as a single species in BPP, and three replicate analyses A11 (joint species delimitation and species tree estimation) were performed using the same parameters as previously mentioned.

### 2.6. Divergence model testing using ABC

Species delimitation analyses perform well to determine species number and species phylogeny, but speciation history may be more complex than a simple dichotomic process as is a species tree. We used the PCA results to identify possible cases of hybridization between groups of individuals. More specifically, the positioning of C4 (Tunisian) individuals in the PCA suggested a possible hybridization event between C3 and C5 (see Results). In addition, the C4 individuals displayed incongruent genetic signals between mitochondrial and nuclear markers previously described (Weber et al., 2015, 2014), as their mitochondrial haplotypes (COI) were closely related to the brooding species C5, whereas their nuclear genotypes (intron i51) were shared with the broadcast spawning species C3. It is noteworthy that i51 was shown to be monomorphic in the brooding species C5 and C6 (Weber et al., 2015).

Then, we used an ABC framework to test eight different models of species divergence (or scenarios) for the *Ophioderma longicauda* species complex, including or excluding hybridization events (Fig. 2). The posterior probability of each scenario, as well as effective sizes, divergence time of each event and admixture rate were estimated using ABC implemented in DIYABC v2.1.0 (Cornuet et al., 2014). Six summary statistics were used to estimate posterior probability of parameters: For the ‘one sample summary statistics’, the number of haplotypes, the number of segregating sites and the mean of pairwise differences were used. For the ‘two-sample summary statistics’, the number of haplotypes, the mean of pairwise differences (between groups) and the F_ST_ statistics (between groups) were used. For ABC analyses, data from the six *Ophioderma longicauda* clusters were used, excluding the outgroups. In addition, nine markers were excluded due to their low amplification success in some clusters. Three sequence groups were defined, each one with a different mutation model. The first group included 19 transcriptome-based markers, the second group included the mitochondrial marker COI and the third group included two introns (Table S2). Default priors were used in preliminary analyses (800,000 simulated datasets) and were then adjusted using posterior distributions and pre-evaluation verifications. When each posterior probability of parameters fell in the prior range in preliminary analyses, 8,000,000 simulated datasets were used to estimate posterior probabilities of parameters and each scenario. Model checking was performed using each available summary statistic, to verify that the parameter values of observed data belonged to posterior distributions.

## 3. Results

### 3.1. Presence of strong incomplete lineage sorting among clusters

Using transcriptome based markers we successfully amplified, sequenced and sorted 30 informative genetic markers (Table S2). Network analyses showed that the majority of markers displayed incomplete lineage sorting, except the mitochondrial marker COI and, although partially, the markers 68241_I.I, i50_II and 98699 (Fig. S1). Not only reciprocal monophyly is not observed among previously-identified species (brooding C5 and broadcast spawning C3 in Crete, (Weber et al., 2017, 2015)) but the large majority of alleles are shared among these species (Fig. S1). K2P distances among mitochondrial lineages ranged from 0.8% between L3 and L3b to 10.7% between L2 and L6 within the *O. longicauda* complex, whereas it ranged from 8.7% between L2 and *O. phoenia* to 11.8% between L6 and *O. phoenia* when considering the three outgroup species (Table S3).

### 3.2. Multivariate analyses identify six genetic clusters

The DAPC showed unambiguously that the optimal number of clusters (i.e. the minimal BIC value) was six (Fig. S2A). The six clusters were very distinct with nearly no overlapping in the 2D representation (Fig. 3A) and 100% probability of memberships for each individual (Fig. S2B). The cluster C1, including all individuals of mitochondrial lineage L6 (from Dakar and Madeira) forms a well-defined group distant from the five other groups (Fig. 3A, Table 1). The cluster C2 includes all L5 individuals from Dakar, whereas the widely distributed broadcast-spawning cluster C3 encompasses all L1 and L5 individuals from Canary Island to Lebanon (Fig. 1, Table 1). The cluster C4 includes all L3b Tunisian individuals (Fig. 1). Finally, the brooding individuals are distributed in two different genetic clusters, incongruent with mitochondrial lineages but congruent with geography. Cluster C5 includes all L2 and L3 individuals sampled in Greece, whereas cluster C6 includes all L2 and L4 individuals sampled in Cyprus and Lebanon (Table 1). The DAPC run without the mitochondrial COI marker provided the same clustering, showing that this marker did not bias the clustering process (Fig. S3). The PCA gave essentially the same results as the DAPC, except that the higher genetic diversity of the broadcast spawners was more visible (Fig. 3B-C). In addition, C4 was even closer to C3 and C5, highlighting the capacity of PCA to explore the natural distribution of individuals and its power to detect potential hybridization events. Furthermore, one individual from Madeira assigned to the cluster C1 was not clustering with the other C1 individuals from Dakar, but was rather at mid distance between the C1 and C3 individuals, suggesting the potential presence of a hybrid between C1 and C3. Yet, further sampling would be required to properly assess this hypothesis. Nevertheless, we excluded this individual from further analyses. Finally, pairwise F_ST_ among clusters were high, ranging from 0.19 between C2 and C3 to 0.47 between C1 and C5 (Table 2). In addition, all F_ST_ values were significant after a permutation test (Table 2).

**Figure 3:**
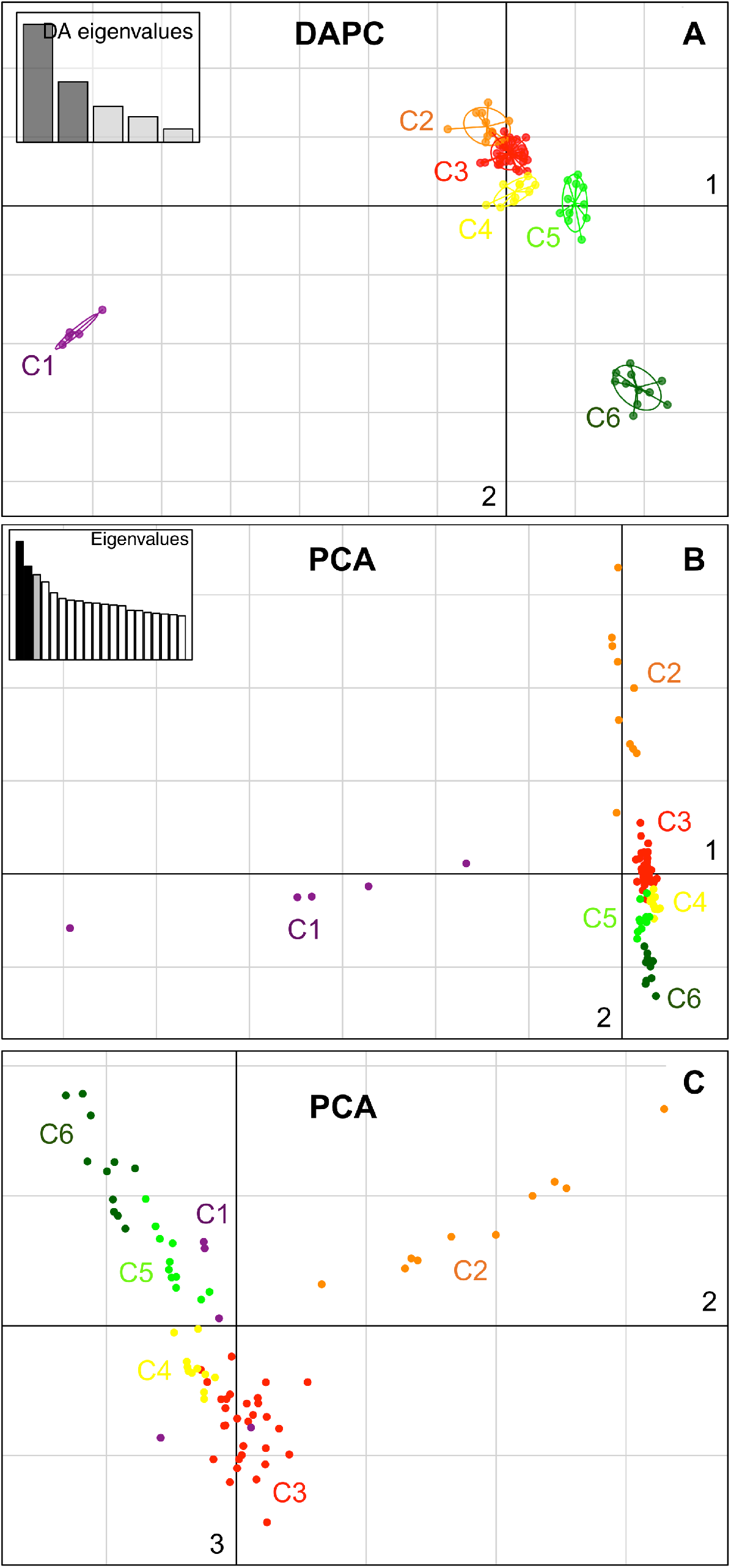
**A**: Results of the Discriminant Analysis of Principal Components (DAPC). The six different genetic clusters are displayed. **B**: Results of the Principal Components Analysis (PCA). Axes 1 and 2 are plotted. Individuals are colored according to their genetic cluster found in DAPC. **C**: Results of the Principal Components Analysis (PCA). Axes 1 and 3 are plotted. Individuals are colored according to their genetic cluster found in DAPC.

**Table 1:**
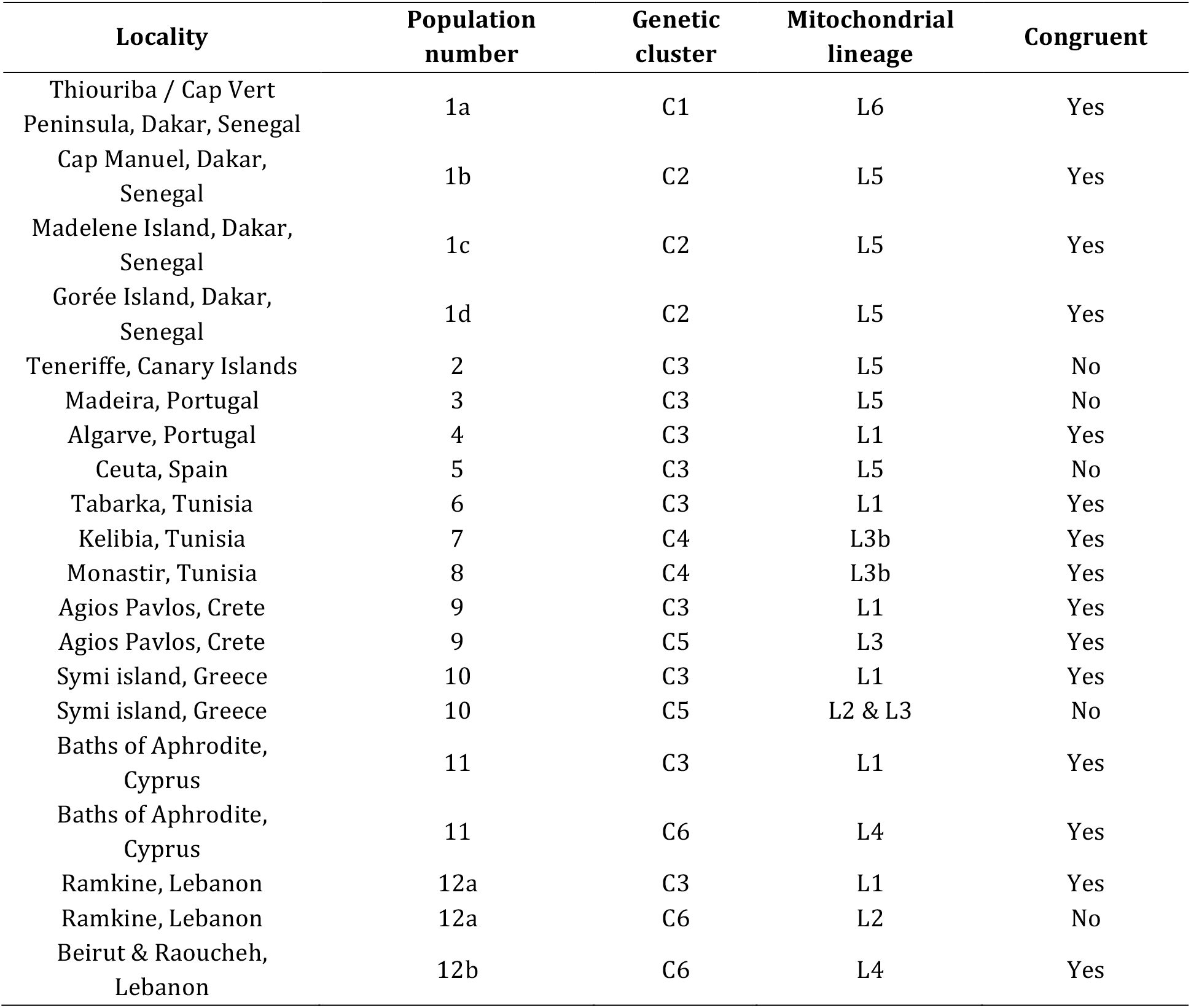
Correspondence between genetic clusters found in DAPC, mitochondrial lineages and sampling locations of individuals used in this study. The population numbers refer to the number indicated in Figure 1. C1-C6: genetic clusters or clusters found in DAPC analysis. L1-L6: *Ophioderma longicauda* mitochondrial lineages defined in Boissin et al., 2011. Congruent: congruence between mitochondrial lineage and nuclear data (genetic cluster).

**Table 2:** F_ST_ values (W&C) among the six genetic clusters based on 30 genetic markers. Significant F_ST_ values after a permutation test (1000 permutations) are highlighted in bold. **: 0.001<P-value<0.01; ***: P-value < 0.001.

| $F_{ST}$ | C2 | C3 | C4 | C5 | C6 |
| --- | --- | --- | --- | --- | --- |
| C1 | **0.28742\*\*\*** | **0.22029\*\*\*** | **0.43463\*\*\*** | **0.47247\*\*\*** | **0.46308\*\*\*** |
| C2 |  | **0.19154\*\*\*** | **0.35779\*\*\*** | **0.37697\*\*\*** | **0.38870\*\*\*** |
| C3 |  |  | **0.20687\*\*\*** | **0.32060\*\*\*** | **0.30226\*\*\*** |
| C4 |  |  |  | **0.39383\*\*\*** | **0.36829\*\*** |
| C5 |  |  |  |  | **0.36176\*\*\*** |

### 3.3. The multispecies coalescent provides divergence time estimates

BPP was used to jointly perform species (or lineage) delimitation and species tree estimation using the six genetic clusters and the three outgroup species. In the three replicate analyses, the six *Ophioderma* genetic clusters C1-C6 and the three outgroups species (*O. teres, O. cinerea* and *O. phoenia*) were fully supported (posterior probability of nine species=1; Table S4). Furthermore, the species tree of the *O. longicauda* species complex was also highly supported (C1 most ancestral, C5 & C6 more recently diverged (Fig. S4); posterior probability=0.92-0.99), although the full topologies were different due to different placements of the three outgroups (Table S4). Using the divergence time of the geminate species *O. teres* and *O. cinerea / O. phoenia* (at least 2.8 mya; (Lessios, 2008)), the divergence times within the *Ophioderma longicauda* species complex were inferred to be at least 537,000 years ago [95% CI: 445,223-682,795] (Table 3; Fig. S4).

It is noteworthy that the BPP analyses run on a sub-dataset of nine C3 individuals and eleven C5 individuals considering each individual as a candidate species in the starting tree gave unsupported results, with unstable numbers of estimated species and low posterior probabilities among replicate analyses (6, 3 and 2 species; Table S5). Therefore, BPP performed poorly to delimit species without a meaningful starting species tree. On the opposite, the DAPC succeeded in finding the true number of species and affecting individuals to them (Fig. S5).

### 3.4. A divergence scenario supports past hybridization

After pre-evaluation of the priors for the eight scenarios (Fig. S6A), posterior probabilities of scenarios tested with ABC indicated that the most probable scenario was the scenario 5, including hybridization between C3 and C5 (PP=0.67; Table S6). The second most likely scenario was the scenario 7, also including a hybridization event between C3 and C5 (PP=0.26; Table S6). The remaining scenarios were not supported. After model checking of scenario 5 (Fig. S6B), parameter estimates indicated that the widespread broadcast spawning cluster C3 displayed the largest effective population size, 3 to 10 times larger than the effective population sizes of the brooding species C4, C5 and C6 (Table 4; Fig. S7). The divergence time estimates indicated that C1 split from other *O. longicauda* clusters about 512,000 generations ago and that the broadcasters and the brooders split about 222,000 generations ago (Table 4). The hybridization event giving rise to C4 was estimated about 90,000 generations ago, with a high proportion of C4 genome originating from C3 (about 86.8%; Table 4). Overall, the divergence events of the *O. longicauda* species complex follow a pattern of West to East differentiation (Fig. 1).

**Table 3:** Posterior probabilities of parameters estimated with BPP. Theta = 4*Ne*μ. Tau = expected number of mutations per site. Minimum divergence times in years are calculated from the minimum divergence time of the geminate species pairs *O. teres* and *O. phoenia/O. cinerea* (2.8 mya, Lessios, 2008). Results for replicate 2 are displayed. See Fig. S4 for the full species tree and the positioning of branch lengths (Tau).

| replicate 2 | mean | median | mode | 2.5% CI | 97.5% CI | mode<br>[years] | 2.5% CI<br>[years] | 97.5% CI<br>[years] |
| --- | --- | --- | --- | --- | --- | --- | --- | --- |
| Theta_C1 | 0.005196151 | 0.005109 | 0.004971 | 0.003570775 | 0.007423225 |  |  |  |
| Theta_C2 | 0.007639499 | 0.007532 | 0.006841 | 0.005636 | 0.01023622 |  |  |  |
| Theta_C3 | 0.01682958 | 0.016739 | 0.014894 | 0.012518 | 0.02186435 |  |  |  |
| Theta_C4 | 0.002738336 | 0.002679 | 0.002408 | 0.0018 | 0.004092 |  |  |  |
| Theta_C5 | 0.001103252 | 0.001069 | 0.001277 | 0.000639775 | 0.001702 |  |  |  |
| Theta_C6 | 0.002360826 | 0.002304 | 0.002319 | 0.001494775 | 0.003635225 |  |  |  |
| Theta_Oteres | 0.004313273 | 0.004187 | 0.004004 | 0.002433 | 0.006873225 |  |  |  |
| Theta_Ophoen | 0.00736498 | 0.007178 | 0.006682 | 0.0046 | 0.01109122 |  |  |  |
| Theta_Ociner | 0.01173528 | 0.011572 | 0.011885 | 0.008069 | 0.016437 |  |  |  |
| Tau_C1.C6 | 0.01596195 | 0.015895 | 0.014907 | 0.013522 | 0.018639 | 8920624 | 8091814 | 11153922 |
| Tau_C2.C6 | 8.09527E-05 | 0.000054 | 0.00001 | 0.00001 | 0.000343 | 5984 | 5984 | 205258 |
| Tau_C3_C4_C5_C6 | 0.000131744 | 0.000113 | 0.000072 | 0.000024 | 0.000325 | 43086 | 14362 | 194486 |
| Tau_C4_C5_C6 | 0.000151832 | 0.000146 | 0.000156 | 0.000037 | 0.0003 | 93353 | 22141 | 179526 |
| Tau_C5_C6 | 0.000152483 | 0.000142 | 0.000155 | 0.00003 | 0.000341 | 92755 | 17953 | 204061 |
| Tau_C1 | 0.000899098 | 0.000883 | 0.000898 | 0.000744 | 0.001141 | 537380 | 445223 | 682795 |
| Tau_C2 | 0.000818136 | 0.000811 | 0.000804 | 0.000675 | 0.000998 | 481128 | 403932 | 597222 |
| Tau_C3 | 0.000686385 | 0.000688 | 0.000721 | 0.000533 | 0.000831 | 431460 | 318957 | 497286 |
| Tau_C4 | 0.000534565 | 0.000535 | 0.000553 | 0.000399 | 0.000685 | 330925 | 238769 | 409917 |
| Tau_C5 | 0.000382091 | 0.00038 | 0.00035 | 0.000247 | 0.000516 | 209446 | 147809 | 308784 |
| Tau_C6 | 0.000382091 | 0.00038 | 0.00035 | 0.000247 | 0.000516 | 209446 | 147809 | 308784 |
| tau_Oteres | 0.004415418 | 0.004374 | 0.004679 | 0.002896 | 0.006119 | 2800000 | 1733020 | 3661723 |
| Tau_Ophoen | 0.004148281 | 0.004135 | 0.003983 | 0.00274 | 0.005535 | 2383501 | 1639667 | 3312246 |
| Tau_Ociner | 0.004148281 | 0.004135 | 0.003983 | 0.00274 | 0.005535 | 2383501 | 1639667 | 3312246 |
| Tau_Ophoen_Ociner | 0.000267135 | 0.000153 | 0.000003 | 0.000005 | 0.001263225 | 1795 | 2992 | 755937 |
| Tau_Ot_Op_Oc | 0.01244562 | 0.0124385 | 0.012251 | 0.0093731 | 0.01559 | 7331225 | 5609036 | 9329344 |

**Table 4:**
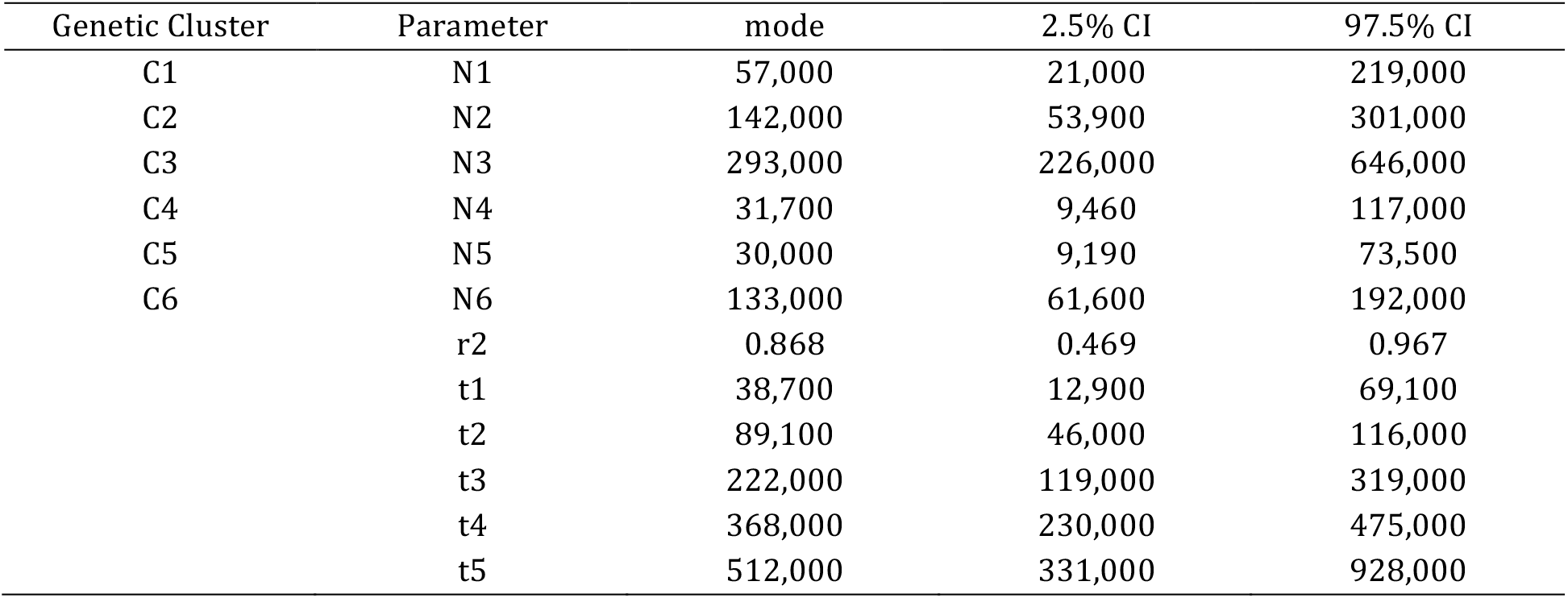
Posterior probability values of estimated parameters for scenario 5 by ABC. N = effective population size; t = divergence times in number of generations, see details in Figure 3. r2 = admixture rate of C3 to C5. q025 and q975 indicate the range of 95% confidence interval.

| Genetic Cluster | Parameter | mode | 2.5% CI | 97.5% CI |
| --- | --- | --- | --- | --- |
| C1 | N1 | 57,000 | 21,000 | 219,000 |
| C2 | N2 | 142,000 | 53,900 | 301,000 |
| C3 | N3 | 293,000 | 226,000 | 646,000 |
| C4 | N4 | 31,700 | 9,460 | 117,000 |
| C5 | N5 | 30,000 | 9,190 | 73,500 |
| C6 | N6 | 133,000 | 61,600 | 192,000 |
|  | r2 | 0.868 | 0.469 | 0.967 |
|  | t1 | 38,700 | 12,900 | 69,100 |
|  | t2 | 89,100 | 46,000 | 116,000 |
|  | t3 | 222,000 | 119,000 | 319,000 |
|  | t4 | 368,000 | 230,000 | 475,000 |
|  | t5 | 512,000 | 331,000 | 928,000 |

## 4. Discussion

### 4.1. Species limits and divergence history deciphered in the O. longicauda species complex

In this study, six *Ophioderma* species (C1-C6) were unambiguously delimited, one of them (C4) most likely originating from the hybridization of C3 and C5. So far, only two *Ophioderma* species have been described in the Eastern Atlantic; *Ophioderma longicauda* (Bruzelius, 1805), from Dakar to Spain in the Atlantic and in the Mediterranean, and *Ophioderma wahlbergii* Müller & Troschel, 1842 in South Africa, even though it was recently shown that the Mediterranean sympatric C3 & C5 and C3 & C6 are different biological species (Weber et al., 2015, 2014). The emergence of the *Ophioderma* genus occurred most likely around the Caribbean Sea, before the closing of the Panama Isthmus. Indeed, most currently recognized extant species (26/28) of this genus thrive in this region (Stöhr et al., 2009) and the oldest confirmed *Ophioderma* fossil (about 10 million years old, Tortonian, early Late Miocene), is from South America (Martínez and Río, 2008). The most divergent species C1 occurring in West Africa split from the common ancestor of C2-C6 at least half a million years ago. Nevertheless, it is possible that the actual divergence time of the species complex is much older as partial closure events of the Panama Isthmus occurred before its final closure around 2.8 mya (e.g. sea catfishes, about 10 mya (Stange et al., 2018)). For the divergence times within *O. longicauda* species complex (C2-C6), we rather refer to the DIYABC estimates, as the divergence model is more accurate.

Given that C1 and C2 were sampled in very close localities (11-17 km apart) and yet genetically very distant, this is further evidence that C1 and C2 are different biological species. Interestingly, Greef (1882) described a new species, *Ophioderma guineensis* Greef, 1882, from West Africa (Gulf of Guinea), which was later considered conspecific with O. *longicauda*, as its distinguishing morphological characters were assumed to fall within the variability of *O. longicauda* (Madsen, 1970). It is possible that this *O. longicauda* “variety” is actually the different biological species that we define here as C1. Yet, fresh samples from this locality are required to test this hypothesis.

Two other broadcast spawners were found, C2 in Senegal and the widespread C3 (from Canary Islands to Lebanon), whereas two species corresponded to brooders (C5 in Greece and C6 in Cyprus and Lebanon). The cluster C4, occurring in Tunisia, is most likely also a brooder, as it displays typical characteristics of brooders (e.g. mitochondrial lineage close to the brooding C5; small effective population size; ecological preference to low depth (Weber et al., 2014)). Gonad examinations of C4 specimens were unsuccessful to determine their reproductive strategy as sampling was performed outside the reproductive season. Yet, it is known that brooders occur in this region as brooding specimens were previously sampled in Tunisia in 1849 and 1924 (Stöhr et al., 2009). Unfortunately, molecular characterization of these samples failed due to poor DNA quality. Interestingly, the most likely origin of the cluster C4 is hybridization between C3 and C5, confirming our initial hypothesis. A formal taxonomic revision of *Ophioderma longicauda* is in progress (Stöhr et al., n.d.). The ancestral strategy of the *Ophioderma* genus is broadcast spawning, as all *Ophioderma* from the Western Atlantic and *O. longicauda* C3 are broadcast spawners. Brooding evolved most likely about 222,000 generations ago in the common ancestor of C5 and C6. Unfortunately, generation time in *O. longicauda* is unknown, although it is known that after they reach sexual maturity, they reproduce once a year (broadcast spawner C3 (Fenaux, 1972); brooders C5 and C6 (Stöhr et al., 2009; Weber et al., 2014)). Interestingly, another independent evolution of brooding occurred in O. *wahlbergii*, which displays much larger and fewer young in its bursae than O. *longicauda* C5 (Landschoff and Griffiths, 2015).

### 4.2. Discrepancy between mitochondrial and nuclear histories cautions the sole use of mitochondrial data for species delimitation

Mitochondrial barcodes such as COI have been widely used in species delimitation and species complex discovery (e.g. (Hebert et al., 2003)) due to the numerous advantages of mitochondrial DNA such as its ubiquity, ease of amplification, high mutation rate and finally its reduced effective size compared to nuclear DNA which makes isolated populations diverge by genetic drift (and eventually reach reciprocal monophyly) more rapidly. In fact, it allowed in the first place the discovery of the *O. longicauda* species complex (Boissin et al., 2011; Stöhr et al., 2009), and it is still efficient to discover additional cryptic species, including brittle stars (Boissin et al., 2017). Nevertheless, mitochondrial lineages did not correspond to species (i.e. genetic clusters) in many cases. For instance, the cluster C3 encompasses individuals displaying the lineages L1 and L5 (4.4% divergence). The same applied for the cluster C5 (lineages L2 and L3, 1.1% divergence) and the cluster C6 (lineages L2 and L4, 2.5% divergence). This is most likely the result of ancient introgression events. Mitochondrial DNA is particularly prone to both selective and introgression sweeps (Currat et al., 2008; Galtier et al., 2009; Pons et al., 2014; Toews and Brelsford, 2012), in contrast to nuclear DNA, and introgression has been proposed in sea urchins (Bronstein et al., 2016). Finally, selection events may be responsible for the retention of particular mitochondrial haplotypes. This study emphasizes the necessity of using nuclear markers to accurately delimit species.

### 4.3. DAPC, BPP and ABC: a potential new combination of methods for species delimitation

Here we used three methods to delimit species in the *Ophioderma longicauda* species complex using 30 genetic markers, of which 25 were transcriptome-based. This is a high number of sequence markers, given that from the 28 studies presented in a review on species delimitation, only two used more than 10 genetic markers (Carstens et al., 2013). The first step was performed using DAPC clustering, based on multi-locus genotypes (e.g. the sequence information was not used, only the allele frequencies) and revealed the presence of six distinct genetic clusters. We showed that this type of clustering approach is powerful in presence of strong incomplete lineage sorting, since diagnostic differences are not needed in order to find genetic clusters, only frequency differences. Therefore, clustering approaches are appropriate for recently diverged species. Then, we verified that these six clusters were confirmed under the MSC framework using BPP that, in addition to providing a tree and estimating the most likely number of lineages, allows the estimation of parameters such as effective population sizes and divergence times. It is noteworthy that the use of BPP alone failed to delimit species on a subset of data. Although BPP has been recently criticized to delimit genetic structure rather than species (Sukumaran and Knowles, 2017), which was not ignored by BPP’s authors (e.g. Rannala 2015), we believe that species delimitation based on genetic data is accurate in the *Ophioderma* case as many genetic clusters are sympatric but do not exchange gene flow (e.g. C1-C2; C3-C4; C3-C5; C3-C6), confirming that they are actual biological species. In addition, different morphological characters were found among C1, C3 and C5 after reanalysis of several specimens (Stöhr et al, *in preparation*). We acknowledge that some uncertainties remain for the allopatric genetic clusters C2-C3 and C5-C6, which might be structured populations from the same species, even though when one compares the genetic distances among these pairs of clusters they are similar to genetic distances among species.

Finally, we propose to go one step further than discovering and delimiting species by inferring a more realistic divergence history, including hybridization, with model testing using ABC. Such methods allow the comparison of complex models including hybridization, reticulate evolution and demographic events (Roux et al., 2013). We found that the most supported scenario included a hybridization event between the broadcast spawners C3 and the brooders C5. Some additional past hybridization events may also have occurred between the divergent C1 and the broadcast spawner C3. Indeed, an individual sampled in Madeira displayed many common alleles with C1, but also many common alleles with C3. Yet, due to the presence of a single potential C1-C3 hybrid, we were not able to test this hypothesis. Nevertheless, this suggests that hybridization and introgression might have been be common in *Ophioderma* species.

A previous study (Camargo et al., 2012) tested the accuracy of BPP (Yang and Rannala, 2010), spedeSTEM (Ence and Carstens, 2011) and ABC methods (Csilléry et al., 2012) for species delimitation. Based on simulations, the authors found that BPP was overall the most accurate, ABC displaying an intermediate accuracy and spedeSTEM the lowest accuracy. All methods displayed lower accuracy when gene flow was incorporated, yet ABC displayed the lowest decrease in accuracy to delimit species. Rather than finding the overall best species delimitation method, we propose to use several consecutive methods to first find the number of distinct genetic entities, and then to estimate the divergence scenarios, therefore taking advantage of the best qualities of each method.

Albeit successful, our pipeline based on exonic amplified markers relies on preexisting genomic resources to develop genetic markers, contrary to other methods such as RAD-sequencing and associated techniques (Davey et al., 2011). RAD-sequencing has been successfully used to delimit species (e.g. (Pante et al., 2015)), yet the efficiency of this method diminished drastically with genetic distance of compared species. Indeed, Pante et al. (2015) report that >70% of loci were lost when species displaying 0.028% of mitochondrial divergence were compared (1-2 myr divergence time) and 97% of loci were lost for species displaying 2.2% of mitochondrial divergence (9-16 myr divergence). This is expected given that the majority of RAD loci are found in non-coding fast evolving DNA. Here, we could successfully retrieve 84-100% of markers for C1-C6 (10.7% maximum mitochondrial divergence (Table S3); divergence at least 537,000 years ago (Table 3)) and 54% of the markers for the outgroup species (11.8% maximum mitochondrial divergence (Table S3); divergence at least 7.3 million years ago (Table 3)), highlighting that our exon-based method performs better than RAD sequencing for distantly related species, but also allows comparing closely related species. In addition, due to their longer sequences compared to RAD loci, our method allows the analysis of haplotype networks (Fig. S1). Therefore, coding sequence markers are useful to compare simultaneously closely and distantly related species. To circumvent the use of individual PCR amplification, one could use our analytic framework with exon-capture data, a method shown efficient to capture exons displaying up to 12% of sequence divergence ((Hancock-Hanser et al., 2013; Hugall et al., 2015) for exon-capture specific to brittle stars). Until now, these data have mainly been used for phylogenomic purposes (O’Hara et al., 2017, 2014) but they could as well be used for cryptic species delimitation with multilocus genotype approaches.

### 4.4. Conclusion

To conclude, the use of three distinct methods with coding sequence markers allowed comparisons at the within- and between-species levels, and bridging the gap between them. We emphasize the power of multilocus genotypes to delimit recently diverged species displaying incomplete lineage sorting and the ability of ABC to uncover the most realistic divergence history of a species complex. We propose that these approaches can be helpful to resolve other complex speciation histories.

## Acknowledgments

We are very grateful to the many people who contributed to sampling of *Ophioderma longicauda* specimens: Helmut Zibrowius, Christos Arvanitidis, Thanos Dailianis, Elena Sarropoulou, Magdalini Christodoulou, Zined Marzouk, Didier Weber, Thi Weber and Philipp Moser. Many thanks to Francisco Alonso Solis-Marin and Harilaos Lessios for providing *Ophioderma* outgroup samples. We also would like to thank Laurent Abi-Rached for his advice on phylogenetic analyses, Arnaud Estoup for his advice on DIYABC and the genomic sequencing platform Genotoul (INRA, Toulouse) for the Illumina sequencing. Finally, we thank the support team of sciCORE (center for scientific computing, University of Basel, http://scicore.unibas.ch/) for providing access to computational resources, especially Pablo Escobar Lopez.

## Funding

A.A-T.W was supported by a scholarship from the French Ministry of High Education, Research and Innovation. S. S. and A. C. did not receive any specific funding for this study.

## Data statement

The raw Miseq reads were deposited on Dryad Digital Repository and are accessible on https://doi.org/10.5061/dryad.5ks03 (The raw data of the present study are from the same Miseq run as for the study: Weber et al., 2015).

